# Persistent Hypersomnia Following Repetitive Mild Experimental Traumatic Brain Injury: Roles of Chronic Stress and Sex Differences

**DOI:** 10.1101/2022.08.03.502603

**Authors:** Edwin Portillo, Xiaomei Zi, Yeonho Kim, Laura B. Tucker, Amanda Fu, Lauren A. Miller, Krystal S. Valenzuela, Genevieve M. Sullivan, Amina K. Gauff, Fengshan Yu, Kryslaine L. Radomski, Joseph T. McCabe, Regina C. Armstrong

## Abstract

Traumatic brain injury (TBI) is often more complicated than a single head injury. An extreme example of this point may be military service members who experience a spectrum of exposures over a prolonged period under stressful conditions. Understanding the effects of complex exposures can support proper evaluation and care for patients experiencing persistent symptoms. We designed a longitudinal series of non-invasive procedures in adult mice to evaluate the effects of prolonged mild exposures. We assessed anxiety, depression, and sleep-wake dysfunction as symptoms that can impact long term outcomes after mild TBI. Unpredictable chronic mild stress (UCMS) was generated from a variable sequence of environmental stressors distributed within each of 21 days. Subsequently, mice received a mild blast combined with closed-head mild TBI on five days at 24-hour intervals. TBI components were either five linear force impacts, or a novel alternating repetitive mild TBI (Ar-mTBI) model of linear and rotational (CHIMERA) impacts over five days to produce diffuse pathology. In males and females, UCMS induced anxiety without depressive behavior. Persistent hypersomnia, specifically increased sleep during the active dark period, was found through 6-12 month time points in male mice receiving UCMS with repetitive blast plus TBI events, or surprisingly after UCMS alone. Sleep-wake dysfunction was not found with TBI events alone and was not found in females under any conditions. These results identify prolonged stress and sex differences as important considerations for sleep-wake dysfunction. Furthermore, this reproducible hypersomnia is similar to excessive daytime sleepiness reported in patients, which may inform treatments.

## INTRODUCTION

Traumatic brain injury (TBI) is heterogeneous in terms of the type and severity of injury, the resulting pathology, and the breadth and timing of symptom expression. This heterogeneity presents significant challenges to evaluation and care. Mild TBI is by far the most common TBI diagnosis and may resolve well over time; yet, for reasons that are not yet clear, a significant proportion of patients diagnosed with mild TBI suffer from persistent symptoms [1, 2]. Indeed, early symptoms of anxiety and/or depression are predictive of worse functional outcomes in patients with mild TBI [3]. Sleep-wake disorders are common after TBI, even in patients diagnosed as mild TBI, which can impair daily functions and contribute to neurodegenerative processes that negatively impact long term brain health [4, 5]. Prolonged stress can also increase the risk for anxiety, depression, and sleep disorders [6]. In patients with mild TBI, co-morbid post-traumatic stress disorder and poor sleep, which are of increased concern for military service members, are each predictive of poor neurobehavioral outcomes [2]. How repetitive head injury events intersect with stressful experiences to influence symptom expression and long term outcomes is not clearly understood, but is important to consider in the diagnosis and treatment of mild TBI.

Complex experiences of repetitive mild TBI with environmental and/or emotional stressors are particularly relevant to military service members, who may experience events of head impact and/or acceleration, blast exposure, and stress due to traumatic experiences, adverse conditions and sleep deficiency. Natural history studies in military and civilian patients with TBI will continue to be important for understanding the heterogeneity of events, including stress and blast exposures, and patient factors such as age and sex/gender that influence symptom expression [2, 7, 8]. To complement these clinical studies in understanding factors contributing to TBI outcomes, specific components of the stress and mild TBI events can be modeled in animals. Exposures can then be tested in permutations to identify the resulting behavioral and pathological responses and work toward effective interventions.

The current studies focus on sleep behavior, which is a clinically meaningful outcome measure for TBI and stress exposures and is also a tractable therapeutic target for improving mental health [9]. Sleep-wake disturbances most commonly associated with TBI are insomnia and hypersomnia/pleiosomnia, sleep-related breathing disorder, circadian rhythm disorder, and parasomnia/movement disorder [5, 10]. Excessive daytime sleepiness is a hypersomnia that occurs during the waking period and is not related to the extent of activity, in contrast to fatigue [5, 10]. TBI significantly increases the likelihood to develop sleep disorders among patients followed out to at least one year post-injury, including veterans diagnosed with TBI [11, 12]. Patients with mild TBI may initially have overlapping sleep problems (e.g. poor sleep quality with excessive daytime sleepiness, or fatigue) but by one year post-injury many patients have one specific sleep problem, which may implicate distinct underlying mechanisms [11].

Several sleep-wake disturbances associated with persistent symptoms in patients after TBI have been replicated in animal models of TBI, including insomnia, excessive daytime sleepiness, and pleiosomnia (Sandsmark and Lim, 2017). Animal models have been particularly useful for identifying neural circuits and molecular mechanisms that promote wakefulness, rapid eye movement (REM) sleep, and non-REM sleep along with control of circadian rhythms [13]. Mouse model studies have revealed critical roles of astrocytes in sleep regulation and the relationship of sleep to neurodegenerative disease processes [14–16]. Continuing to improve our understanding of the neural circuits and mechanisms underlying sleep homeostasis and pathology in animal models is important to inform clinical decisions in the use of current treatments and to identify novel therapeutic targets or interventions for specific sleep-wake disturbances.

We developed a longitudinal study design to test the contribution of specific stress and head injury components to neurobehavioral symptom expression and sleep-wake dysfunction. We modified the use of established mouse models of unpredictable chronic mild stress (UCMS) [17], repetitive mild blast exposure [18], and repetitive closed-head concussive linear and rotational mild TBI models [19, 20]. The study design is for each component (environmental stressors, repetitive blast and mild brain injury, or isolation stress) to be mild so as not to independently produce significant symptoms but rather to result in persistent symptom expression when experienced in combination over a prolonged period of time. In studies of mild TBI, significant differences have been observed in outcomes between men and women [7, 8]. Therefore, sex was evaluated for the response to stress and head injury exposures. The longitudinal analysis of anxiety and sleep patterns between events across time reveals remarkable effects of stress and sex on the induction of sleep-wake dysfunction that models excessive daytime sleepiness in humans and persists at chronic time points of up to one year.

## METHODS

### Mice

All mice were treated in accordance with guidelines of the Uniformed Services University of the Health Sciences and the National Institutes of Health Guide for the Care and Use of Laboratory Animals. C57BL/6J (IMSR Cat# JAX:000664, RRID:IMSR_JAX:000664) 7 week old male and female mice were purchased from Jackson Laboratories. Except as required for experimental procedures, mice were socially housed in 35 cm x 16.5 cm x 18 cm enrichment cages (2-5 mice per cage), exposed to 12:12 hour light-dark cycle, at controlled room temperature, and with free access to food and water. All experimental procedures were conducted during the daytime light cycle (06:00-18:00) after the mice had acclimated for at least 3 days in their home cage.

### Study Design

Cohorts of 24 mice were randomly assigned to groups for sham (n = 12) or experimental (n = 12) procedures. Male and female mice were run in separate cohorts to avoid intersex influences on behavior. A pre-determined series of exposures was conducted with combinations of 21 days of unpredictable chronic mild stress (UCMS), 5 days of blast with impact TBI, and 30 days of single housing as an isolation stress. For each experiment of 24 mice, these components were run in the same sequence and followed the same timeline to maintain equivalent age comparisons when components were omitted. Longitudinal assessments for anxiety and depression included the elevated zero maze, tail suspension test, and social interaction, while some cohorts also included the sucrose preference test. Longitudinal assessment of sleep behavior was conducted using the non-invasive PiezoSleep mouse behavioral tracking system.

### Stressor Procedures and Behavioral Assessments for Anxiety, Depression, and Sleep-Wake Profile

As in prior reports of UCMS [17], a series of mild stressors was selected for exposures in an unpredictable pattern over 21 days (Table 1). Testing revealed that mice did not avoid areas of a test cage where predatory odor solution was applied (data not shown). Therefore, predatory odor was omitted as a stressor. See Supplemental Materials for detailed procedures of the stressors shown in Table 1 and behavioral assessments for elevated plus maze, tail suspension test, sucrose preference test, social interaction, and sleep-wake function.

**Table 1:** Schedule of environmental stressors in unpredictable chronic mild stress (UCMS) protocol

|  | MORNING |  | AFTERNOON | OVERNIGHT |
| --- | --- | --- | --- | --- |
| day 1 | cage change/weigh | foot shock | restraint | strobe |
| day 2 | weigh | orbital shaker | forced bath | cage tilt |
| day 3 | weigh | forced bath | restraint | no bedding |
| day 4 | weigh | forced bath | foot shock | cage tilt |
| day 6 |  | no procedure |  |  |
| day 7 |  | sleep disruption 8AM-8PM |  |  |
| day 8 | cage change/weigh | foot shock | orbital shaker | strobe |
| day 9 | weigh | restraint + shaking | forced bath | no bedding |
| day 10 | weigh | restraint | forced bath | wet bedding |
| day 11 | weigh | foot shock | restraint | strobe |
| day 12 | weigh | orbital shaker | foot shock | cage tilt |
| day 13 |  | no procedure |  |  |
| day 14 |  | sleep disruption 8AM-8PM |  |  |
| day 15 | cage change/weigh | forced bath | orbital shaker | no bedding |
| day 16 | weigh | foot shock | restraint + shaking | cage tilt |
| day 17 | weigh | restraint | orbital shaker | wet bedding |
| day 18 | weigh | foot shock | forced bath | cage tilt |
| day 19 | weigh | restraint + shaking | forced bath | strobe |
| day 20 |  | no procedure |  |  |
| day 21 |  | sleep disruption (continuously for 12 hrs starting 6:00-8:00 AM) |  |  |

Details of each stressor procedure are provided in the Supplemental Materials

### Repetitive Blast (r-Blast) Procedures

At 8-9 weeks of age, mice were exposed to five simulated blast exposures delivered at 24 hour intervals in an Advanced Blast Simulator (ABS) (ORA, Inc., Fredericksburg, VA), as previously detailed [18]. ABS procedures were performed with mice in the prone position for frontal blast exposure. Each mouse was anesthetized with 3% isoflurane and the head and body was secured to a wooden tongue depressor, wrapped in gauze, placed in mesh to prevent rotational movement, and positioned to face the oncoming blast as previously detailed [21, 22]. Sham mice were anesthetized and followed the same procedures except that the blast was not deployed. Across the cohorts in the current experiments, the blast exposure mean peak incident overpressure was 21.08 ± 0.8 psi, which is consistent with 21.7 ± 1.11 psi exposures in our prior characterization of the blast pathology [18]. Each blast exposure was immediately followed by a closed head impact, so that the blast and impact combination was completed within 10 min.

### Closed Head Repetitive Mild TBI (r-mTBI) from Linear Impact

After the ABS exposure on each of the five days, an Impact One Stereotaxic Impactor (Leica Biosystems; Buffalo Grove, IL) was used to produce the linear r-mTBI concussive impacts, as previously detailed [19]. A series of five impacts separated by 24-hr intervals was chosen to target the period of decreased glucose uptake observed 24 hrs after a single mild TBI in rats [23]. Mice were anesthetized with 2.0% isoflurane in O_2_ and then the hair was shaved and depilated with Nair. Mice were positioned in a stereotaxic frame with the ear bars inverted and covered with rubber stoppers to maintain the head in a fixed position. A 3-mm flat tip with rounded edges was used to impact the scalp approximately over bregma (velocity set at 4.0 m/sec; depth of 1.0 mm; dwell time of 200 ms). Sham mice underwent identical procedures to the r-mTBI mice without receiving impacts. Body temperature was maintained with a warming pad. After each impact procedure, the time for the mouse to right itself from the supine to prone position was recorded to quantify the righting reflex as a surrogate measure of loss of consciousness.

### Alternating Repetitive Mild TBI (Ar-mTBI) from Linear and Rotational Impact

To generate more diffuse pathology without increasing the impact severity, we took advantage of two different mild closed head TBI models. The repetitive linear force r-mTBI model (above) is a fixed head impact onto the scalp over bregma that has been characterized for five impacts at 24 hour intervals [19]. The Closed-Head Impact Model of Engineered Rotational Acceleration (CHIMERA) allows for head movement after mild impacts onto the scalp over bregma and has been characterized using five impacts at 24 hour intervals [24]. We tested this rotational injury over five days using an impact energy of 0.5J [24] and found only subtle axon damage beyond the frontal cortex (data not shown) and so increased to 0.6J for the current studies, which is still in the range of prior mild TBI studies with the CHIMERA device [25]. The Ar-mTBI model used the linear impact device for TBI on days 1, 3, and 5 with rotational impact on days 2 and 4 using the CHIMERA device. Sham mice underwent identical procedures to the Ar-mTBI mice without receiving impacts. Body temperature was maintained with a warming pad. The righting reflex times were recorded after each combination of blast followed by linear or rotational impact.

### PiezoSleep Analysis of Sleep-Wake Function

Sleep-wake data were collected using a non-invasive automated scoring in the PiezoSleep system home cages (Signal Solutions LLC, Lexington, KY). Mice were single housed and maintained on a standard 12 hour cycle of daytime light (6:00–18:00) with continuous data collection. A cage floor matt with piezoelectric sensors recorded 4 second epochs and used the 2-4 Hz breathing rhythm of mice to classify intervals of 30 seconds or more as asleep or awake [26]. This piezoelectric sensor system compares well with sleep data collected by visual observation and with electrophysiological discrimination of sleep/wake intervals, yet avoids the surgical procedures of electrophysiological techniques that could confound other assessments of TBI [27, 28].

### Neuropathology

At the completion of the experimental protocol mice were perfused with 4% paraformaldehyde, post-fixed overnight, cryoprotected with sucrose and embedded for cryostat sectioning of the fixed frozen tissues. Sectioning and staining were performed by FD Neurotechnologies (Columbia, MD). Sagittal sections (40 μm) were stained for neurodegeneration with FD NeuroSilver^TM^ Kit. Sagittal sections (14 μm) were stained for hematoxylin and eosin or immunostained for glial fibrillary acidic protein (GFAP; rat-anti-GFAP; #13-0300 Thermo Fisher Scientific; RRID:AB_2532994), ionizing calcium binding adaptor protein 1 (IBA1; rabbit anti-IBA1; #019-19741 WAKO; RRID:AB_839504) and myelin basic protein (MBP; rat anti-MBP; #MAB386, Millipore; RRID:AB_94975).

### Statistical Analysis

The number of mice of each sex that were used to generate the results for each experiment is shown in Table 2. The statistical analysis for each result is explained at the end of each figure legend. Significance was set as p < 0.05. Automated data collection was used in all behavioral assessments. However, weights of mice and sucrose/water solutions require manual data collection.

**Table 2:** Mouse number and sex for data shown in experiments. Parenthesis = died before completion of experiment.

| Experiment Cohorts | Shams | Experimental | Sex |
| --- | --- | --- | --- |
| Figure 1 | 10 (2) | 11 (1) | male |
| Figure 2 | 12 | 11 (1) | male |
| Figure 3, 4 | 6 | 6 | male |
|  | 6 | 6 | female |
| Figure 5, 7 | 12 | 12 | male |
| Figure 6, 7 | 12 | 11 (1) | female |
| Figure 8 | 12 | 12 | male |
| Figure 9 | 11 (1) | 12 | male |
|  | 12 | 12 | female |
| Totals | 93 | 93 |  |

## RESULTS

### Repeated exposure to mild environmental stress in an unpredictable series produces weight loss and anxiety

We first established a protocol of unpredictable chronic mild stressors (Table 1) and a subsequent period of isolation that comprised the stress components of the model (Figure 1A). Body weight was monitored as a sign of overall animal health. Weight loss was observed during the 21-day sequence of varied stress exposures and again during the isolation period (Figure 1B). Mice began to recover weight while still experiencing the UCMS protocol and during the isolation housing, which indicates adaptation of the mice to these mild stress components. Mice exposed to the UCMS protocol and the isolation period exhibited signs of anxiety-like behavior, as indicated by reduced time in the open arm of the elevated plus maze (Figure 1C). The stress components did induce social impairment, which is a function associated with a depressive phenotype in mice [29]. Social interaction was tested for social approach (Figure 1D) based on interaction time exploring an unfamiliar mouse relative to an empty carrier and for social memory (Figure 1E) based on preference for exploring the first now familiar mouse relative to a subsequently introduced unfamiliar mouse. Sham and stressed mice exhibited similar significant preferences in each stage of social behavior testing. Stress exposed mice were not different from shams in preferring pleasurable sucrose-containing water to plain water, which is a test of anhedonia (data not shown). Mice of both conditions also had a similar time interval to immobilization on the tail suspension test of despair or learned helplessness (data not shown). These results indicate that the stress components of the protocol produce weight loss and anxiety without signs of a more severe depressive phenotype.

**Figure 1.**
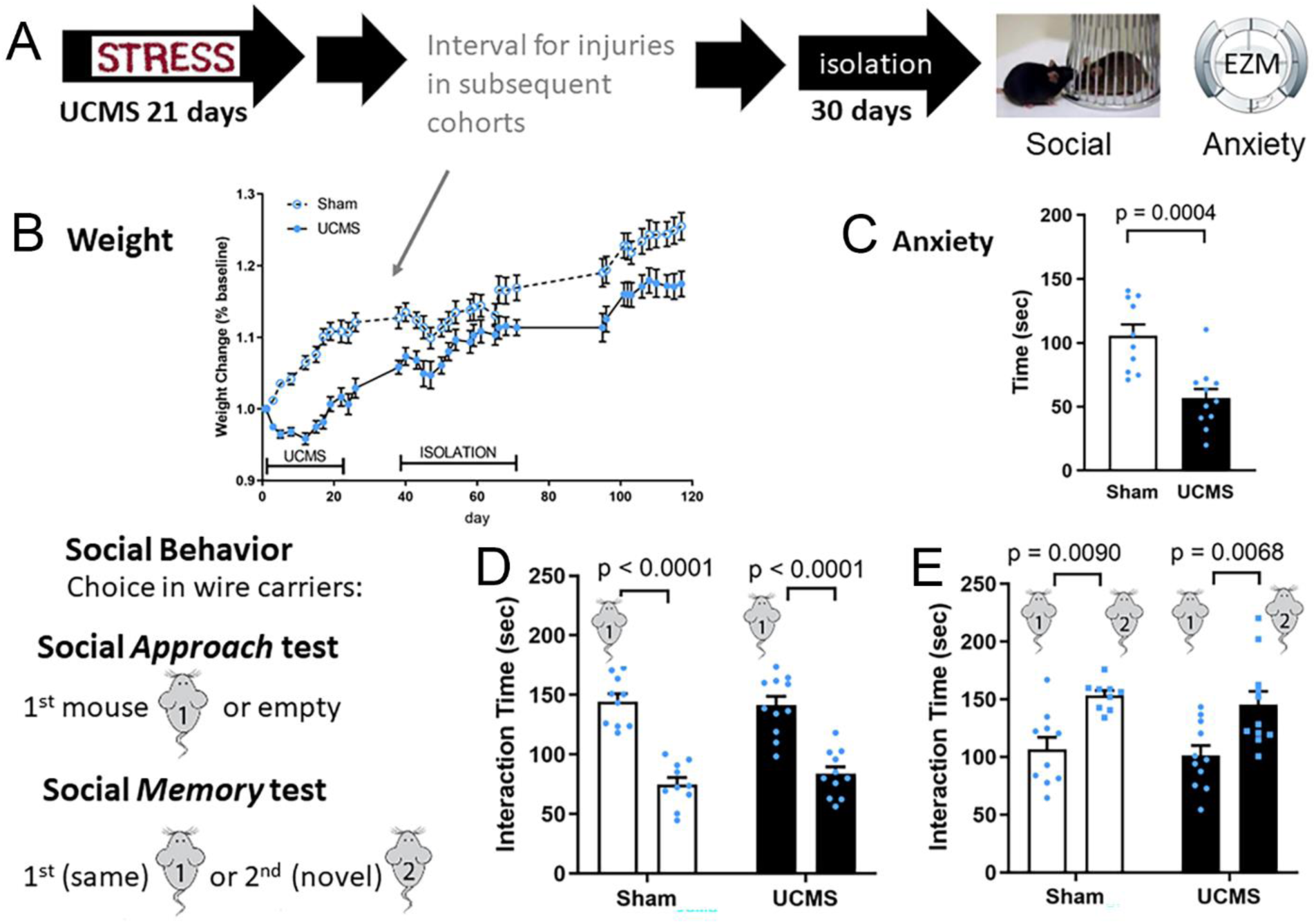
Environmental stress over 21 days induces a mild yet persistent anxiety-like behavior. **A:** Experimental design using male mice (blue symbols). **B:** The mice exhibit weight loss during the 21-day protocol of unpredictable chronic mild stress (UCMS). **C:** The UCMS mice exhibit anxiety-like behavior as indicated by reduced time spent in the open area of the elevated zero maze (EZM), as compared to the sham control mice. **D-E:** Social behavior is not impaired in the UCMS mice. Sham and UCMS mice respond socially to approach an unknown congenic mouse (D), and subsequently remember that first mouse to exhibit more interest in a second novel congenic mouse (E). Statistical analysis with Student’s t-test for EZM and two-way ANOVA with Tukey’s multiple comparison test for social behavior. The number of experimental mice is shown in Table 2. Values shown are mean ± sem.

### Repeated events of unpredictable mild stress and blast exposure with impact TBI result in anxiety and chronic sleep-wake dysfunction

Blast and impact brain injury exposures were integrated between the mild stress components, i.e. UCMS and isolation, to establish a full spectrum of events that resulted in persistent symptoms (Figure 2A). UCMS resulted in anxiety behavior of less time spent in the open arm (Figure 2B). Repetitive injury of a blast exposure (r-blast) followed immediately by a closed head linear impact (r-linear TBI) was performed at 24-hour intervals on 5 consecutive days. The righting reflex time had a longer delay after the head impacts, which is an indicator for loss of consciousness (Figure 2C). Over the 5 days, the righting reflex delay shortened and was not significantly different from the sham group by days 4 and 5. Anxiety behavior persisted after the repetitive injury series (Figure 2D), and then attenuated after the isolation period based on testing at 1, 6, or 12 months post-injury (Figure 2E-G). The mice exposed to stress and head injuries did not exhibit impaired social behavior, anhedonia in the sucrose preference test, or altered immobility during the tail suspension test (data not shown).

**Figure 2.**
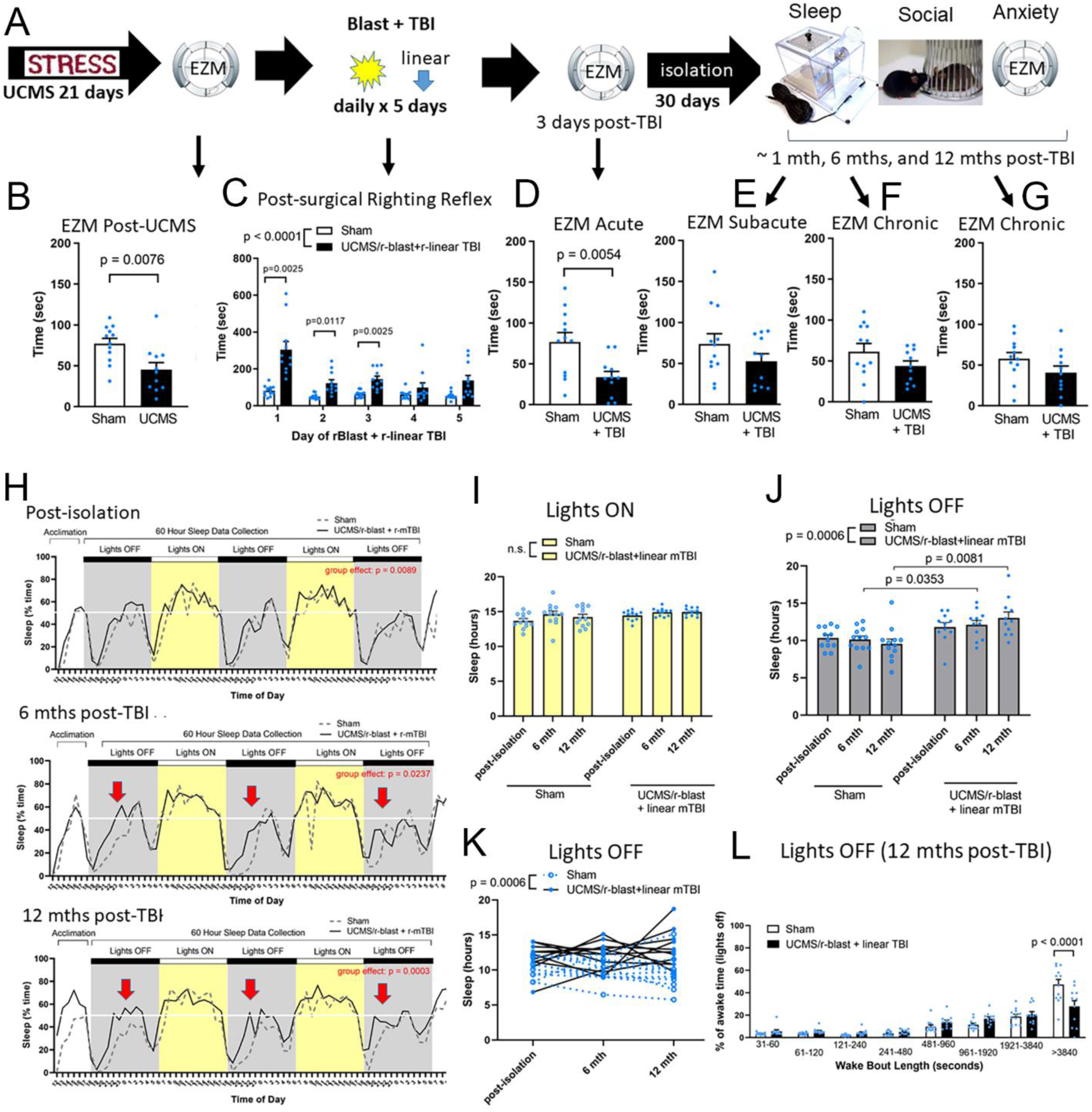
UCMS with repetitive blast and linear TBI results in early anxiety and chronic stage sleep-wake dysfunction. **A:** Experimental design using male mice (blue symbols). **B:** Mice exposed to UCMS exhibit anxiety-like behavior, as indicated by reduced time spent in the open area of the elevated zero maze (EZM) as compared to the sham control mice. **C:** Mice exposed to combined blast combined with linear impact mild TBI (r-mTBI) exhibit a delayed righting reflex period on days 1-3 with this mild repetitive injury protocol. **D-G:** Anxiety effects persist after repetitive head injury (D) and then attenuate to no longer be significantly different from shams after the isolation period (E) and at longer time points of 6 months (F) and 12 months (G) post-TBI. **H:** Mice that experience UCMS with repetitive blast and mild TBI (UCMS/r-blast+r-mTBI) have a significantly different sleep-wake pattern as compared to sham that is most notable during the dark period, which is the awake cycle for nocturnal mice. **I:** Mice of both conditions sleep for similar amount of time when the lights are on. **J-K:** The mice with stress and head injury exposures sleep for an excessive amount of time during the dark (awake) period (J) which can continue across time in individual mice (K). **L:** In the UCMS/r-blast + linear mTBI mice, the percentage of time spent in long wake bouts is reduced as compared to sham mice. Statistical analysis with Student’s t-test for EZM and two-way ANOVA for longitudinal studies of righting reflex time and sleep time, with Holm-Sidak’s multiple comparison test. The number of experimental mice is shown in Table 2. Values shown are mean ± sem.

Sleep-wake behavior was examined as a potential long term effect of TBI that impacts daily function and may contribute to neurodegeneration [5]. Sleep-wake epochs were quantified non-invasively using piezoelectric mats in a home cage over a continuous 72 hour period. Longitudinal studies showed that mice exposed to the stress and brain injury (r-blast + r-linear TBI) protocol had a significantly different sleep pattern from shams at the time points after isolation and 6 and 12 months post-injury (Figure 2H). The sleep time was not altered during the daytime periods with the lights on so that insomnia was not evident (Figure 2I). In contrast, during the normally active overnight dark periods, mice exposed to the stress and brain injury protocol slept for a significantly greater amount of time, which increased through the 12 month time point (Figure 2J, K). The increased sleep corresponded with fragmentation of the long periods of waking without otherwise altering the relative distribution of wake bout lengths (Figure 2L). Therefore, the stress and brain injury protocol produced a pattern of sleep-wake dysfunction with hypersomnia that appears to model excessive daytime sleepiness in patients who cannot maintain wakefulness after TBI [10].

### Diffuse mild pathology is produced by repetitive blast and TBI from alternating linear and rotational models

To better capture the heterogeneity of repetitive mild TBI events, we conducted the same experimental protocol in a novel model of alternating linear and rotational mild TBI (Ar-mTBI). We expected that alternating linear and rotational models over 5 days would achieve diffuse axon damage and pathology without requiring an increase of force for each impact (Figure 3). In combination with the blast exposures, this more diffuse pathology could have effects on the neural circuits between the cortex and ventral brain regions that regulate sleep-wake behavior [13]. At 3 days after the last of the blast plus impact injuries, no overt neuropathology is visible based on histology (Figure 3A), which is consistent with maintaining a mild level of brain injury. Neuroinflammation after Ar-mTBI is evident based on immunostaining of astrocytes (Figure 3B) and microglia (Figure 3C). Myelin staining does not indicate focal lesions but the density of myelination appears reduced in areas of prefrontal cortex and dispersed ventral brain regions (Figure 3D). Silver staining detects degenerating axons in broad areas (Figure 3E, 4A-D). The cerebral cortex under the impact site shows evidence of degenerating neuron cell bodies and axons (Figures 3E, 4A). Descending cortical axons show degeneration down to the gray matter junction with subcortical white matter, which is a hallmark site of axon pathology in human TBI (Figure 4A). Degenerating axons within white matter tracts are visible in the corpus callosum, optic tract, and cerebellum (Figure 4A, B, C). The optic tract had the most extensive axon damage (Figure 4B), and corresponding with this visual system pathology, silver staining was present in the superior colliculus (Figure 4D). This optic tract damage has been well described in reports of CHIMERA pathology [25] and has been associated with optic canal damage in a weight drop model of TBI in mice [30]. Astrogliosis indicated a secondary injury response that quantitatively corroborated pathology in the areas with axon degeneration indicated by silver staining (Figure 4G).

**Figure 3.**
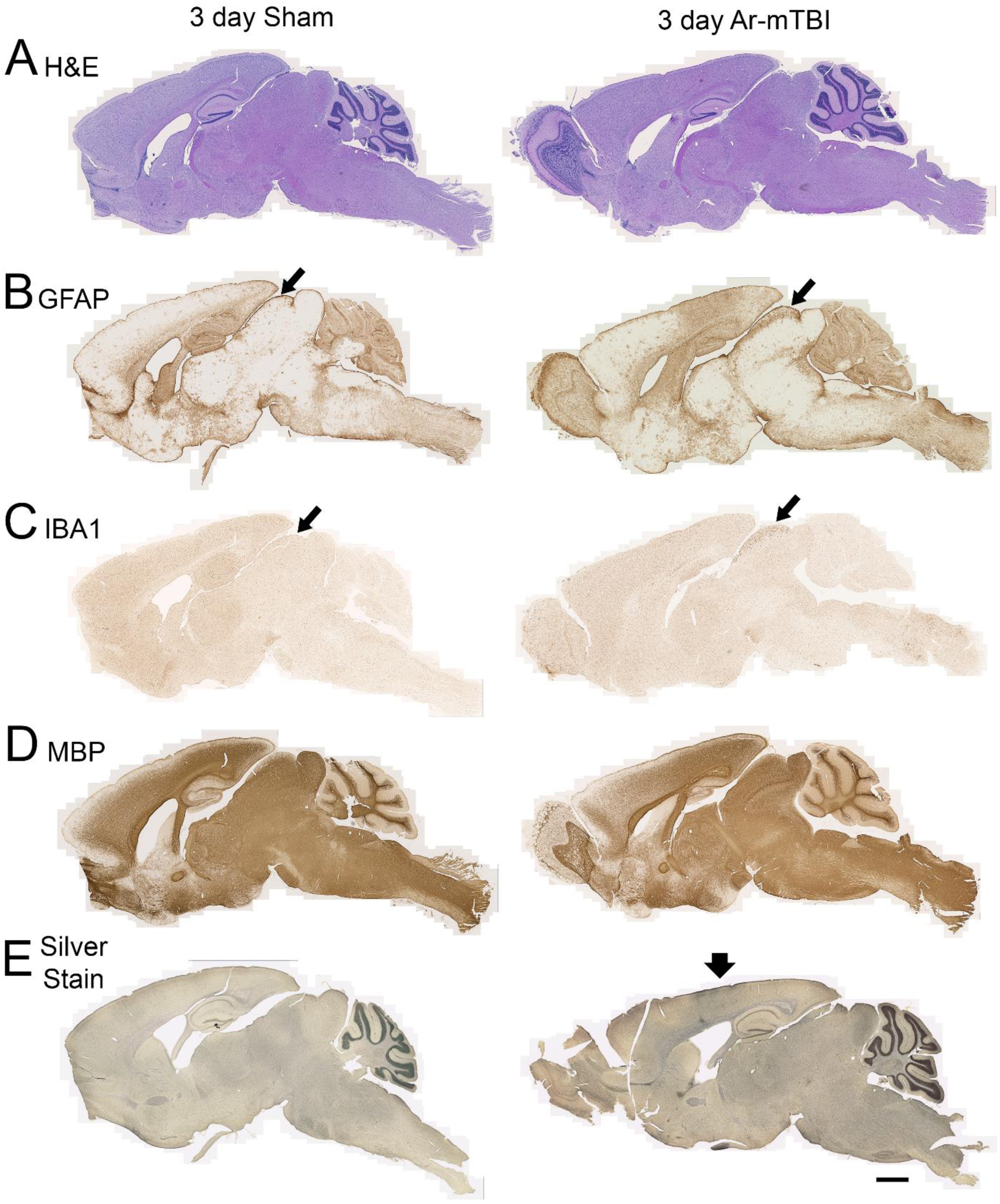
An alternating model of repetitive mild TBI (Ar-mTBI), with blast exposures, produces diffuse pathology. Sagittal sections stained to detect a range of pathological features. Examples shown are sham (left side) and blast with Ar-mTBI at 3 days after the last injury procedure (right side). **A:** Hematoxylin and eosin (H&E) histological staining shows that repetitive mild exposures to blast plus linear or rotational impact did not produce overt pathology, such as cavitation or hemorrhages. **B:** Increased immunoreactivity for glial fibrillary acidic protein (GFAP) indicates astrogliosis after injuries (right) as compared to sham procedures (left). Thin arrows point to the superior colliculus (SC), which shows the most marked astrogliosis. **C:** Immunoreactivity for ionizing calcium binding adaptor molecule 1 (IBA1) indicates a microglia reaction in the SC (thin arrows). **D:** Immunostaining for myelin basic protein (MBP) shows axonal myelination. **E:** Silver staining detects degenerating axons in the cerebral cortex (thick arrow) in the area under the impact site onto the skull at bregma. Degenerating axons are also evident in the optic tract, and to a lesser degree in distributed regions. Examples shown are from female mice with similar findings in male mice. Scale bar is equal to 1 mm (bottom right).

**Figure 4.**
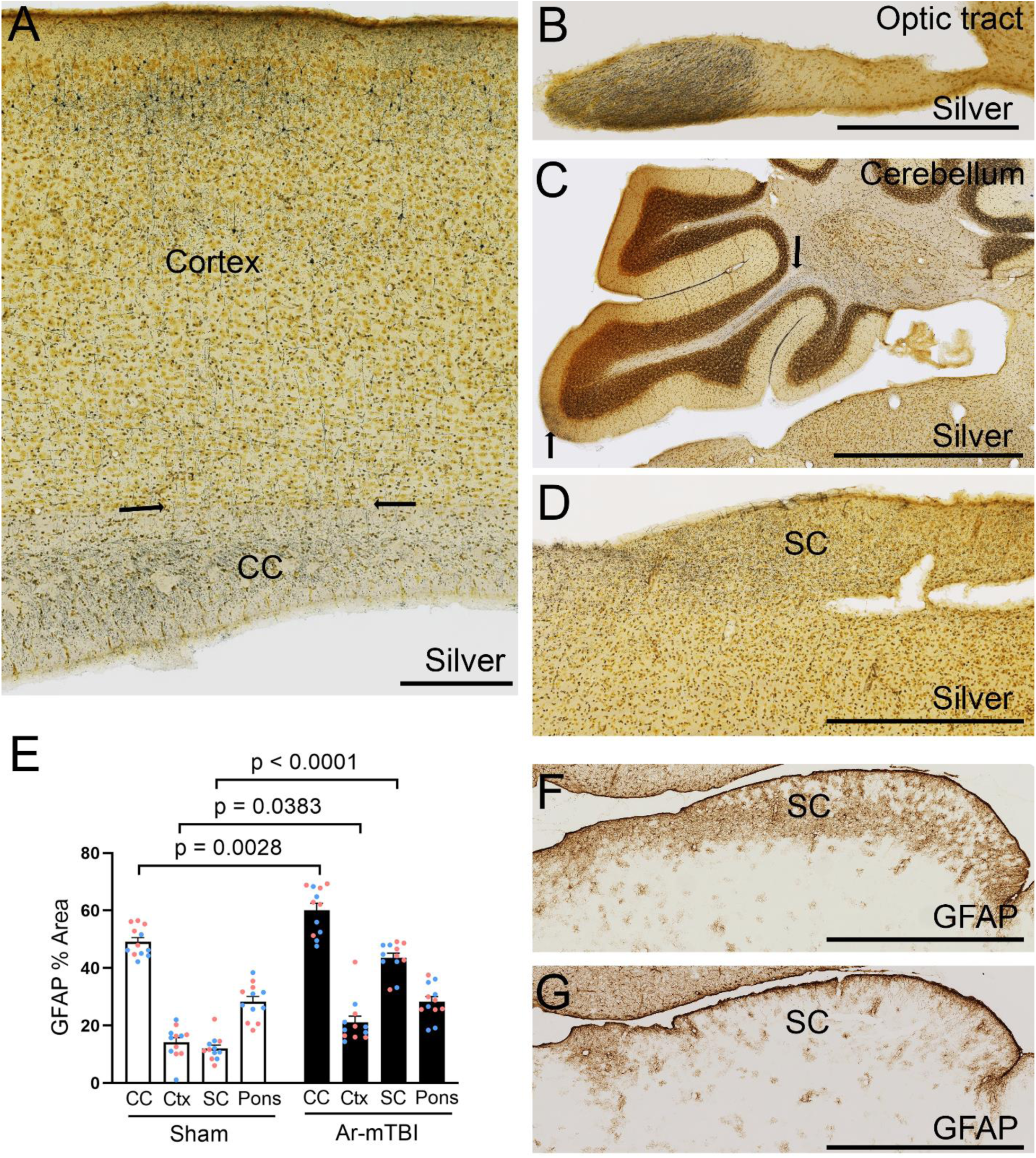
Axon degeneration and astrogliosis are found in diffuse brain regions at 3 days after Ar-mTBI with blast exposures. **A-D:** Silver staining to detect axon degeneration in the sagittal plane. Under the impact site, silver staining labels cortical neurons and axons descending through the gray-white matter junction (between arrows) into the corpus callosum (CC), as well as labeling transverse and longitudinal axons within the CC (A). The optic tract exhibits the most marked axon degeneration after injuries (B). Silver staining of degenerating axon is visible in cerebellar axons in white matter of the arbor vitae (C, upper arrow) extending into the uvula (lobule 9) and at the posterior tip of the lobule (C, lower arrow) in the molecular layer, which is mainly comprised of parallel fibers. Degenerating axons are dispersed within the superior colliculus (SC)(D). **E-G:** Increased GFAP immunolabeling demonstrates significant astrogliosis in the cortex (Ctx) under the impact site, in the CC, and in the SC (E), with the most marked difference found in the SC after injury (F) as compared to sham procedures. (G). The pons region was also quantified as a region that was more remote from the injuries. The pons did not have a significant difference in GFAP immunolabeling (E). Data includes equal numbers of males (blue symbols) and females (pink symbols) Two-way repeated measures ANOVA with Holm-Sidak’s multiple comparison test. The number of experimental mice is shown in Table 2. Values shown are mean ± sem. Scale bars: A = 200 μm, B = 500 μm, C-G = 1 mm.

### Repeated events of unpredictable mild stress and blast exposure with alternating linear and rotational TBI reproduce anxiety and sleep-wake dysfunction

The prior experimental protocol (Figure 2A) was repeated with the Ar-mTBI model to test for reproducibility and/or differences due to inclusion of the rotational model (Figure 5A). This comparison demonstrated that these results were robust and reproducible. The protocol with the Ar-mTBI model produced consistent differences in the righting reflex across each of the 5 days (Figure 5C). UCMS resulted in anxiety behavior (Figure 5B) that persisted after the injury phase (Figure 5D) and then attenuated at longer times post-injury out to 12 months (Figure 5E-G), which is the same time course of anxiety behavior as found using the r-mTBI linear impacts (Figure 2). Sleep-wake dysfunction was evident particularly after the UCMS procedures (Figure 5H). Mice of both conditions slept a similar amount of time during the typical sleep period with the lights on, indicating an absence of insomnia (Figure 5I). Hypersomnia was found during the dark phase after stress and head injury (Figure 5J), indicating that the ability to maintain wakefulness was impaired. Hypersomnia in the dark phase was observed at 12 months post-injury in this protocol with the Ar-mTBI model, similar to the result with the linear r-mTBI model (Figure 2J, 5J). However, this hypersomnia was found at the earlier time points examined in the protocol with this experiment with the Ar-mTBI model (Figure 5J).

**Figure 5.**
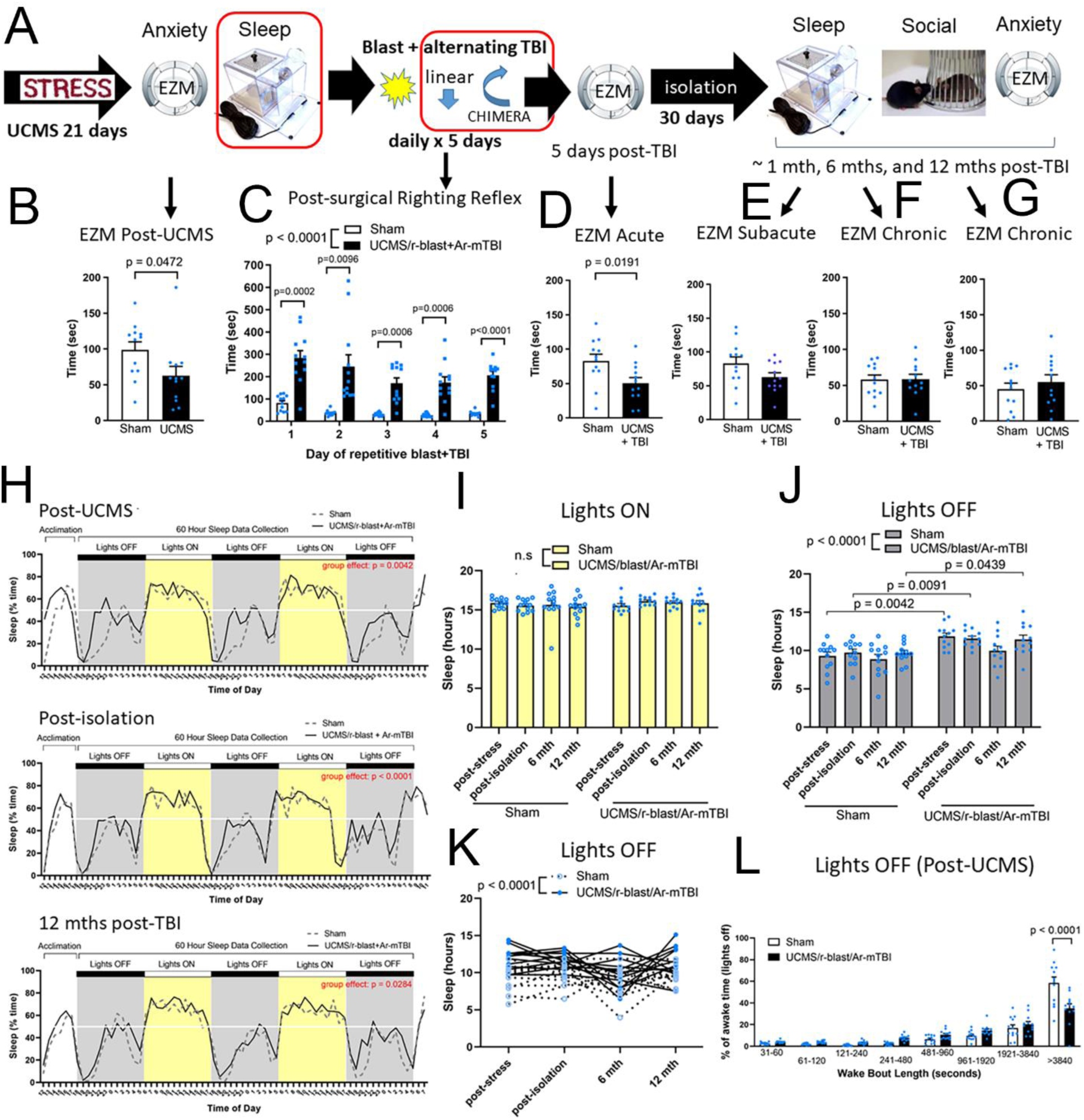
UCMS with repetitive blast and Ar-mTBI produces early anxiety and sleep-wake dysfunction. **A:** Experimental design using male mice (blue symbols) is matched to Figure 2, with two changes. The first sleep studies occur at the end of the UCMS exposures. The post-blast daily injuries alternate impact devices for linear (days 1, 3, 5) and rotational (days 2, 4) forces. **B:** Mice display anxiety-like behavior on the elevated zero maze (EZM) after UCMS, as compared to the sham control mice. **C:** Mice exposed to combined blast and alternating mild TBI exhibit a delayed righting reflex period on days 1-5 with this mild repetitive injury protocol. **D-G:** Anxiety effects persist after repetitive head injury (D) and then attenuate to no longer be significantly different from shams after the isolation period (E) and at longer time points of 6 months (F) and 12 months (G) post-TBI. **H:** Mice that experience UCMS with repetitive blast and alternating mild TBI (UCMS/r-blast+Ar-mTBI) have a significantly different sleep-wake pattern that is most notable during the dark (awake) period. **I:** Mice of both conditions sleep for similar amount of time when the lights are on. **J-K:** The mice with stress and head injury exposures sleep for an excessive amount of time during the dark (awake) period (J) which can continue across time in individual mice (K). **L:** In the UCMS/r-blast+Ar-mTBI mice, the percentage of time spent in long wake bouts is reduced as compared to sham mice. Statistical analysis with Student’s t-test for EZM and two-way ANOVA for longitudinal studies of righting reflex time and sleep time, with Holm-Sidak’s multiple comparison test. The number of experimental mice is shown in Table 2. Values shown are mean ± sem.

Additionally, in the experiments of the Ar-mTBI protocol (Figure 5A), the longitudinal design included a sleep study at the end of the UCMS that was highly informative. Immediately after the UCMS component, the mice display sleep-wake dysfunction with increased sleep time only during the dark period (Figure 5H-K). This finding suggests that UCMS alone is sufficient to impair the ability to maintain wakefulness. The long wake bouts were significantly reduced after UCMS, indicating fragmentation of the waking periods (Figure 5L) which continued to be present during the sleep analysis after the injury and isolation period (data not shown).

### Sleep-wake dysfunction is a sex-dependent effect after repeated events of unpredictable mild stress and blast exposure with alternating linear and rotational TBI

We ran matched studies in female mice (Figure 6) for comparison with the male mice (Figure 5) and found a remarkable sex difference in sleep-wake behavior, yet the anxiety response was equivalent in both sexes (Figure 7). In comparison to sham females, females exposed to UCMS and blast with Ar-mTBI exhibited anxiety behavior and righting reflex effects (Figure 6A-G) as in males (Figure 5A-G). Surprisingly, the female mice did not exhibit disrupted sleep-wake behavior during either the light or dark phase at any time point examined (Figure 6H-J). In addition, in female mice the wake bout lengths were similar in sham mice and those that experienced stress and injuries (Figure 6L). Thus, females did not exhibit impaired wakefulness (Figure 6L), which is in contrast to the findings in males after UCMS (Figure 5L).

**Figure 6.**
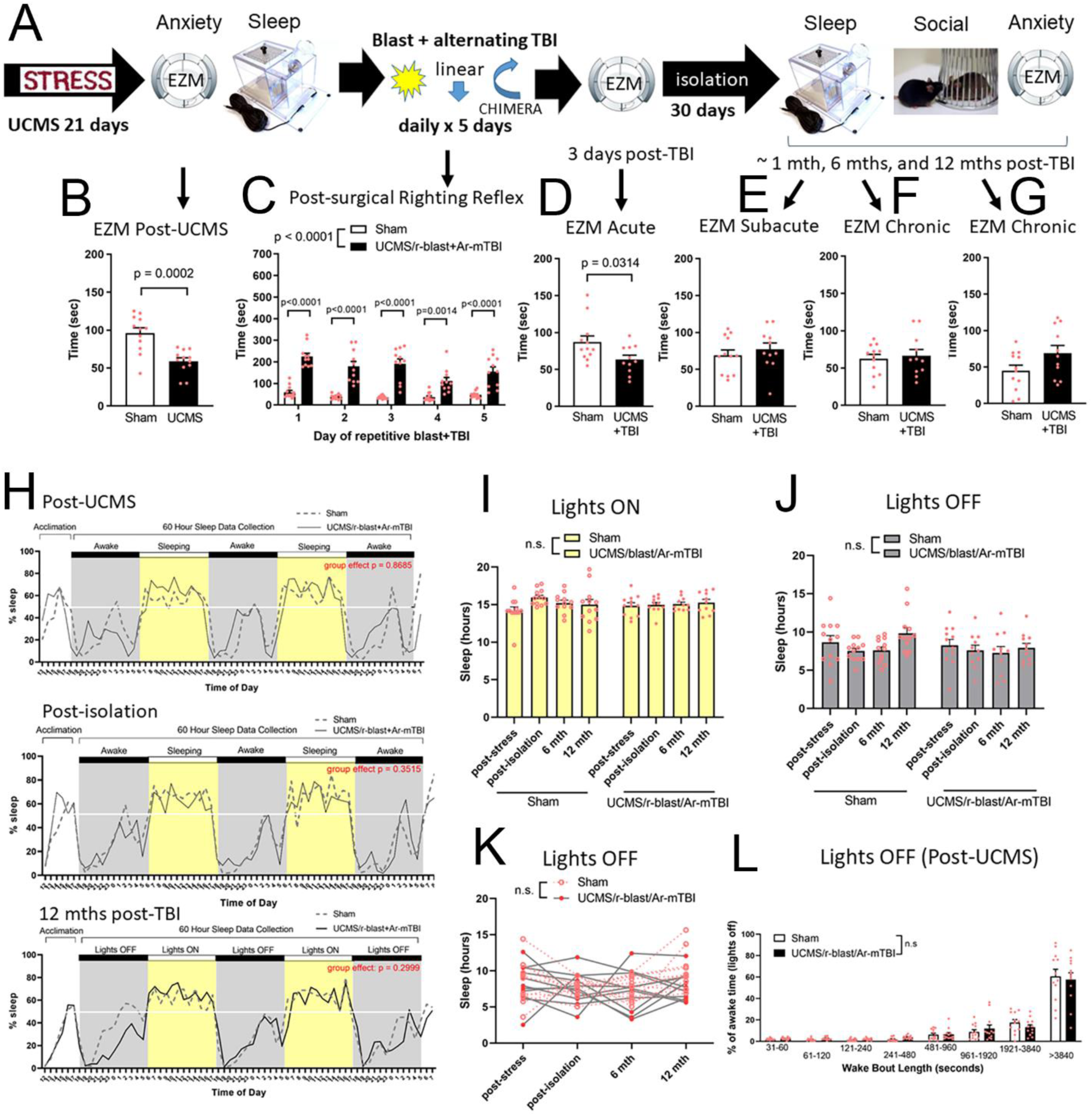
In female mice, UCMS with repetitive blast and Ar-mTBI produces early anxiety, yet does not result in sleep-wake dysfunction. **A:** Experimental design is matched to Figure 5 with the sex changed to female mice (pink symbols). Longitudinal sleep studies are initiated after the UCMS. The post-blast daily injuries alternate impact devices for linear (days 1, 3, 5) and rotational (days 2, 4) forces. **B:** Female mice exhibit anxiety-like behavior on the elevated zero maze (EZM) after UCMS, as compared to the sham control mice. **C:** Female mice exposed to combined blast and alternating mild TBI exhibit a delayed righting reflex period on days 1-5 with this mild repetitive injury protocol. **D-G:** UCMS mice continue to display anxiety behavior after repetitive head injury (D) but this difference from sham mice is attenuated after the isolation period (E) and at longer time points of 6 months (F) and 12 months (G) post-TBI. **H:** Female mice that experience UCMS with repetitive blast and alternating mild TBI (UCMS/r-blast+Ar-mTBI) do not exhibit sleep-wake dysfunction, as compared to sham females. **I-K:** Female mice of both conditions have similar sleep time when the lights are on (I) or off (J, K). **L:** The UCMS/r-blast+Ar-mTBI mice have a similar distribution of wake bouts after UCMS as compared to sham mice. Statistical analysis with Student’s t-test for EZM and two-way ANOVA for longitudinal studies of righting reflex time and sleep time, with Holm-Sidak’s multiple comparison test. The number of experimental mice is shown in Table 2. Values shown are mean ± sem.

This sex difference effect was more directly tested by statistical comparison of the data from males (Figure 5) and females (Figure 6). Female mice had a shorter righting reflex delay after the head injuries, but were not significantly different from the males on any individual days (Figure 7A). For anxiety behavior, the sham mice show a main effect of time for both males and females but do not have a significant difference for sex (Figure 7B). Anxiety behavior of mice exposed to stress and injuries does not show a sex difference, as both sexes have reduced time in the open arm that is maintained without an effect of time (Figure 7C). Importantly, the sleep studies show that males exposed to stress and head injuries exhibit significantly increased sleep during the dark phase at all time points when compared to females (Figure 7D). The sleep time of the individual male and female mice further illustrates the consistent sex difference found with repeated testing that persists for over a year (Figure 7E). The sham males also have a slight increase of sleep during the dark phase but this effect was not robust across time points (Figure 7F). After experiencing the stress and head injury with the Ar-mTBI model, neither male nor female mice exhibited impaired social motivation or social memory, anhedonia in the sucrose preference test, or altered immobility during the tail suspension test (data not shown).

**Figure 7.**
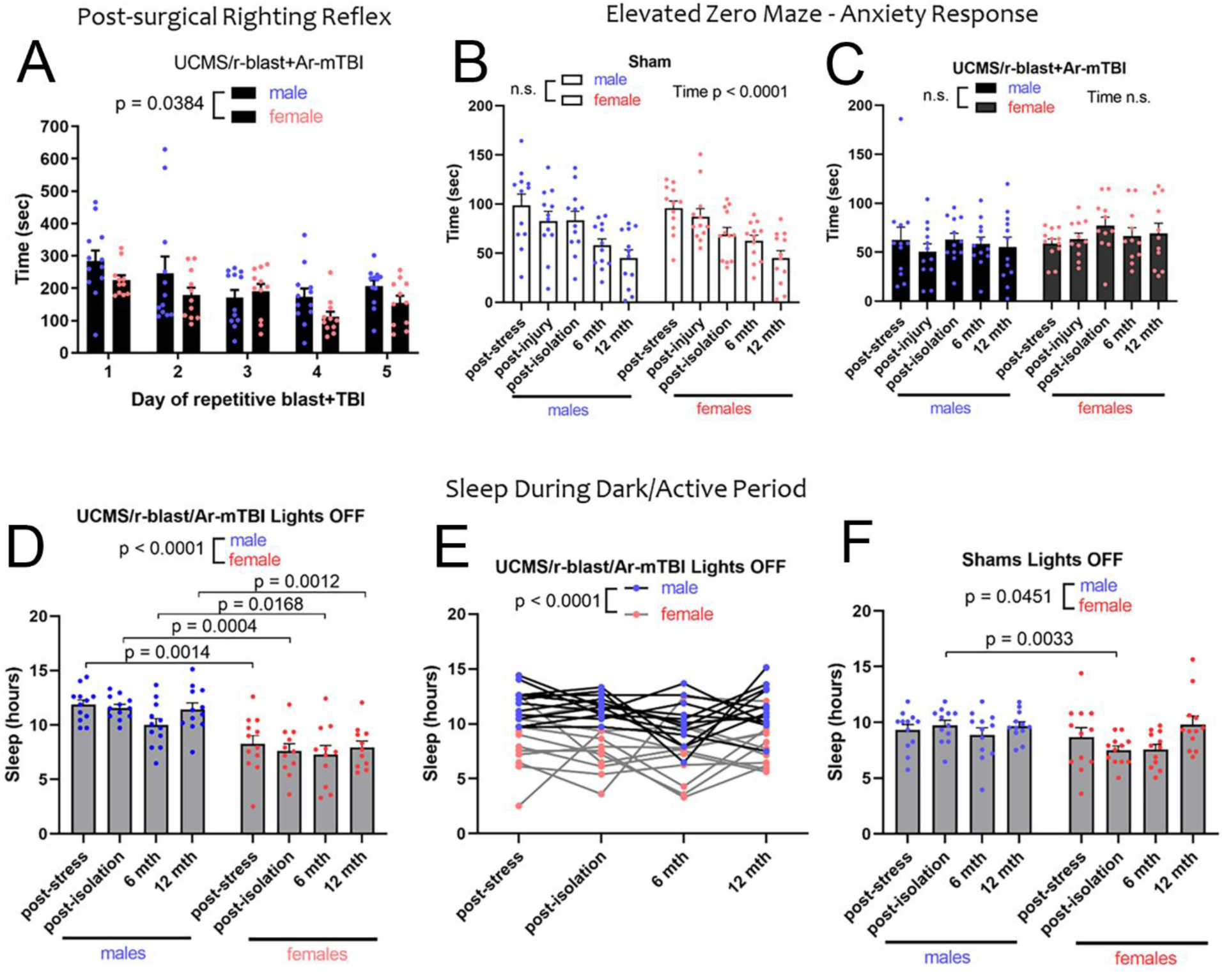
Sex is a significant variable in sleep-wake dysfunction. Data for male (blue symbols) and female (pink symbols) mice extracted from Figures 5 and 6, respectively, for statistical comparison of the effect of sex. **A:** The righting reflex has a main effect of sex but does not a significant effect of sex on individual days when corrected for multiple comparisons. **B-C:** Time spent in the open area of the elevated zero maze indicates a significant main effect of time in sham mice (B) but no effect of sex in sham (B) or mice that experienced the full protocol (C). **D-F:** During the lights off period (gray bars), sleep time is significantly increased in males as compared females, indicating an effect of sex on maintaining wakefulness after stress and repetitive blast with TBI, in the group analysis (D) that is also evident in the individual mouse data (E). Sham mice show a significant, but less robust, effect of sex on sleep time during the dark phase (F). Two-way ANOVA for longitudinal studies of righting reflex time, elevated zero maze, and sleep time, with Holm-Sidak’s multiple comparison test. The number of experimental mice is shown in Table 3. Values shown are mean ± sem.

Overall, these findings demonstrate a profound and prolonged sex-difference effect on sleep-wake behavior in mice that experience a series of stress and head injury exposures. Specifically, male mice exhibit impaired ability to maintain wakefulness, while sleep is not disrupted in female mice.

### Repetitive mild closed head injury does not produce sleep-wake dysfunction in the absence of stress

Next, we separately tested components of the protocol to identify exposures that contributed to hypersomnia in male mice, which revealed that the repetitive blast and head injury component does not produce anxiety or sleep-wake dysfunction (Figure 8). The experimental protocol (Figure 8A) was age- and sex-matched to the experimental design shown in Figure 5 that employed 5 daily head injuries (combined blast and Ar-mTBI impact). Both stress components (UCMS and isolation) were omitted from the experimental design. In place of UCMS, the mice were group housed for the 21 days before head injuries. Similarly, in place of the isolation period, the mice were group housed for 30 days. The mild repetitive head injury procedures, in the absence of stress components, did not produce signs of anxiety (Figure 8B-E) or sleep-wake dysfunction (Figure 8F-G), in comparison to sham mice.

**Figure 8.**
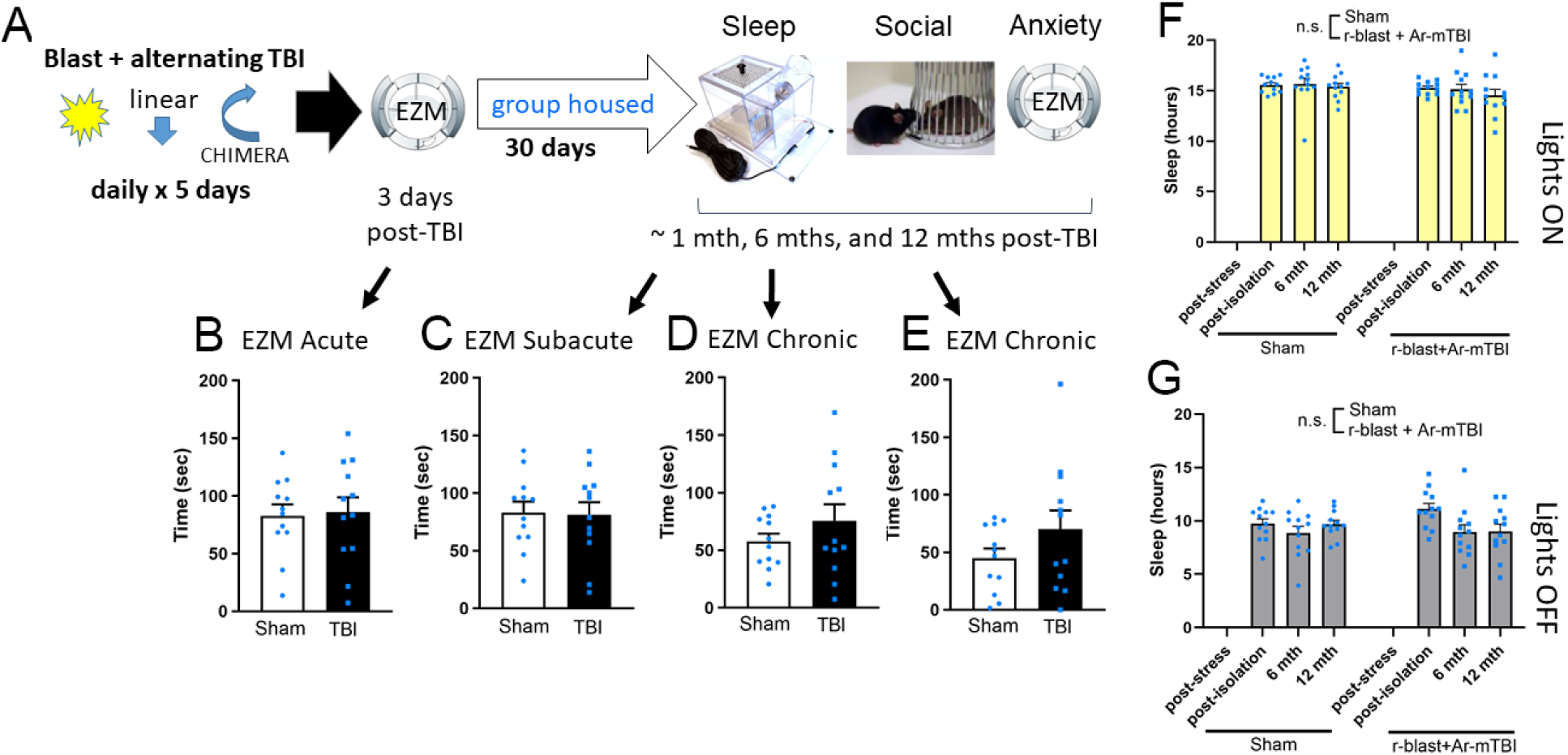
Repetitive blast with Ar-mTBI does not produce anxiety or sleep-wake dysfunction. **A:** Experimental design in male mice (blue symbols) is matched to Figure 5 with omission of UCMS and isolation stress components. The mice are group housed for the same number of days that match each stress component period. The post-blast daily injuries alternating impact devices for linear (days 1, 3, 5) and rotational (days 2, 4) forces exhibited a delayed righting reflex period in injured mice on each day, as compared to sham control mice (data not shown). **B-E:** Mice do not exhibit anxiety-like behavior on the elevated zero maze (EZM) at any time points. **F-G:** Mice that experience repetitive blast and alternating mild TBI (r-blast+Ar-mTBI) have similar sleep time to compared to shams when the lights are on (F) or off (G). Statistical analysis with Student’s t-test for EZM and two-way ANOVA for longitudinal studies of sleep time, with Holm-Sidak’s multiple comparison test. The number of experimental mice is shown in Table 2. Values shown are mean ± sem.

### Stress produces sleep-wake dysfunction in males in the absence of head injury

Finally, we demonstrated that the stress components of the experimental design, in the absence of blast or impact head injuries, were sufficient to impair wakefulness in male mice (Figure 9A). The UCMS component alone significantly increased sleep during the dark phase, as compared to sham mice, and this hypersomnia persisted after the isolation stress period (Figure 9B-C). To test the effect of the stress components with a neurobehavioral assessment indicative of a depressive phenotype, we again examined social behavior and again did not find social approach (Figure 9D) or social memory deficits (Figure 9E).

**Figure 9.**
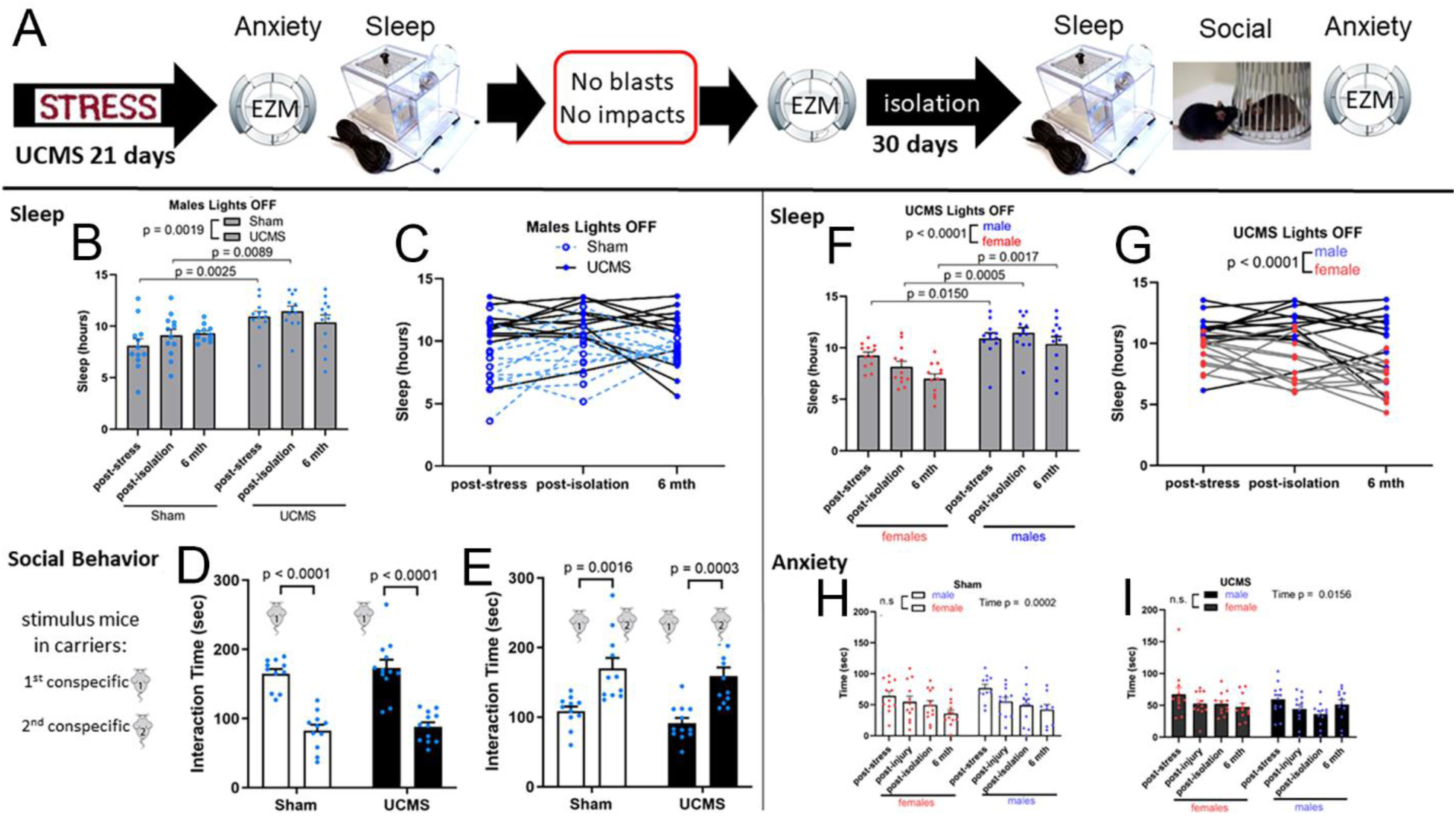
UCMS and isolation stress components produce early anxiety and sleep-wake dysfunction in males. **A:** Experimental design using male (blue symbols) and female (pink symbols) mice is matched to Figure 5 and Figure 6, respectively, with omission of the blast and impact TBI components. **B-C:** In male mice, as compared to shams, mice exposed to stress components exhibit increased sleep during the dark (active) period both at the group (B) and individual level (C). **D-E:** At the 6 month time point, the stress exposures does not impact social behavior in male mice. Stressed and sham mice exhibit significant social approach behavior (D) in response to an initial unfamiliar mouse (1) and a significant social memory response (E) in choosing to explore a novel mouse (2) relative to the initial mouse (1). **F-G:** Male mice exposed to stress components exhibit increased sleep during the dark (active) period both at the group (F) and individual level (G), as compared to females. **H-I:** Male and female mice exhibit similar levels of anxiety behavior on the elevated zero maze (EZM) for both sham mice (H) and those that experience UCMS and isolation stress (I). Two-way ANOVA for sleep and anxiety longitudinal studies with Holm-Sidak’s multiple comparison test or with Tukey’s multiple comparison test for social behavior. The number of experimental mice is shown in Table 2. Values shown are mean ± sem.

An independent cohort of female mice was run in parallel with the male mice following the same experimental design (Figure 9A). These experiments reproduced the robust sex-difference in hypersomnia at each time point (Figure 9F-G) as in our prior comparison of male and female mice (Figure 7D-E), although now the effect was replicated with exposure to only the stress components. The sleep studies show that males exposed to the stress components exhibit significantly increased sleep during the dark phase at all time points when compared to females (Figure 9F). The graphs of the individual male and female mice illustrate the consistent sex difference found with repeated testing that persists through the longest time tested, which was over 6 months (Figure 9G). UCMS also induces anxiety behavior, in comparison to shams that appeared to be similar in male and female mice (Figure 9H-I) in the face of disparate effects on sleep behavior.

This experimental separation of the stress components from the head injury components revealed that the mild but unpredictable and prolonged environmental stress exposures can have a long term effect on the ability to maintain wakefulness. Furthermore, males are more likely to exhibit this effect of environmental stress, even in the absence of head injury events.

## DISCUSSION

The combined results of this study identify chronic stress and sex as significant modifiers of sleep behavior, which is a high priority concern among persistent long term symptoms after TBI. Our UCMS protocol is comprised of relatively mild experiences yet is sufficient to elicit anxiety and impair wakefulness. Impaired ability to maintain wakefulness after UCMS was present in two experiments with male mice (Figures 5 and 9) and absent in two experiments with female mice (Figures 6 and 9). Thus, UCMS reproducibly resulted in anxiety and impaired wakefulness in males while adding repetitive blast with mild TBI, from either of two TBI models (Figures 2, 5), did not produce additional sleep-wake symptoms. Yet, this surprising finding of a hypersomnia effect in male mice that requires UCMS exposure is not fully explained by the anxiety induced in response to UCMS, since males and females exhibited a similar anxiety response.

This hypersomnia during the awake or active period in our male mice is particularly interesting in light of clinical studies reporting that anxiety, but not depression, is associated with excessive daytime sleepiness after TBI [31]. The UCMS protocol produced anxiety behavior at the post-UCMS and post-injury time points but the significant difference from shams was attenuated at the longer time points (Figures 2, 5, 6). However, this attenuation was effectively due to an effect of time in the sham mice (Figures 7B-C, 9G-H), which is consistent with prior results in naïve mice with repeated testing over several weeks [32]. In the current studies, the longitudinal effect on the anxiety response with repeated testing over the one year time frame may also involve an effect of aging in the sham group that is not seen in the mice that are already compromised by the stress and repetitive head injury exposures.

The current experimental design induced anxiety without evidence of depressive signs that have been differentiated based on social interaction in mice [29]. Social function is an important factor in mood disorders after TBI [33]. Importantly, social function was not impaired across any cohorts when tested at multiple time points (see Figures 1, 9). However, social deficits were previously observed in male mice after repetitive mild TBI at 8 weeks of age with social approach deficits found in studies using the linear model [19] but only social memory deficits were found with the rotational model [24]. The lack of social deficits in the current experiments could be due to several factors. In the current protocols, the mild TBI was paired with a preceding blast exposure and/or with the flanking UCMS and isolation stress periods. In addition, the repetitive TBI series was delayed until 12 weeks of age in the current experiments in order to accommodate the preceding UCMS protocol. Interestingly, young age at the time of injury may relate to greater vulnerability to poor social outcomes in patients and in experimental TBI models [34].

Sleep-wake dysfunction can be stressful and sleep fragmentation after TBI may delay or limit recovery [35]. In our experiments, the 21-day UCMS and subsequent 5-day injury procedures were conducted during the daytime with the lights on, which disturbed the main sleep period for mice. Importantly, mice did not exhibit compensatory sleep during the typical lights on sleep phase immediately following the UCMS procedures (Figures 5, 6). Mice exhibited increased sleep time only during the active dark period and did so well into the chronic stages at 6 and/or 12 month time points, which indicates that this hypersomnia is not due to compensation for early sleep disruption during the UCMS or injury procedures (Figures 5, 7, 9).

Mouse models have greatly facilitated characterization of neural circuits controlling sleep-wake function that extend between neurons of prefrontal cortex and multiple ventral brain regions [13]. Particularly relevant to our current findings of increased sleep during the waking period after UCMS (Figures 5, 9) are elegant neural network studies that examined the response to stress and effects on sleep behavior [36]. These studies in male mice identified a midbrain circuit that responded to social defeat stress and induced sleep. More specifically, cells of the ventral tegmental area (VTA) that connected to the lateral hypothalamus elicited non-REM sleep when activated by optogenetic stimulation. Stimulation of the VTA neurons also reduced the stress-induced anxiety. Similarly, network connections between prefrontal cortex and the basolateral amygdala have been studied as models of social dysfunction in autism [37]. The absence of social behavior deficits after UCMS in our experimental design (Figures 1, 9) may be an important clue as to network connections involving the VTA in the anxiety response. Interestingly, chronic social defeat stress and neural network analysis in male mice demonstrated that the VTA to basolateral amygdala dopamine circuit selectively controls anxiety-like behavior but not depression-like behaviors [29].

Analysis of sleep-wake behavior after TBI in mouse models has produced mixed findings, which may be due to heterogeneity of the TBI. Our studies did not find sleep-wake disturbance when the mice experience only the repetitive mild head injuries (Figure 8). However, more severe TBI models in male mice and rats have reported increased sleep and/or increased sleep-wake fragmentation at early times post-injury [10, 38]. Mild fluid percussion injury in male mice caused a persistent inability to maintain wakefulness and decreased orexin neuron activation during wakefulness [39]. Further studies indicate that this impaired wakefulness may result from damage to glutamatergic neurons that synapse onto orexin neurons in the lateral hypothalamus and was mitigated by dietary therapy [39, 40].

The current results are an example in which a single early stage assessment would not have identified the involvement of sex differences in another outcome measure. Specifically, the results on the elevated zero maze indicated similar anxiety behavior in males and females, while only males exhibited sleep-wake dysfunction (Figures 5, 6, 7, 9). These findings support best practices of testing for sex differences on each outcome measure before deciding whether the data should be combined in subsequent experiments. This sex-difference in the outcomes is also intriguing in that, as yet unknown, host factors must also have a role in the interaction of the neural circuits that control anxiety and sleep behavior.

There are several limitations of this study to consider when interpreting the results. The non-invasive piezoelectric analysis of sleep-wake function was selected for the advantages of a enabling a longitudinal study design and avoiding surgical procedures, which could confound the mild stress and injury procedures. However, this sleep data is limited to sleep-wake decisions for sleep time and bout length. More detailed analysis of the sleep-wake architecture using EEG/EMG may inform interpretation of the hypersomnia observed. For example, analysis of sleep-wake microarchitecture can detect differences in the power of delta and theta frequency bands which identified effects of controlled cortical impact TBI in male mice [41]. Furthermore, our experiments compared the effects using two TBI models and showed that the sleep effects did not require rotational force or even head injury, so the TBI pathology was not a focus in the current study. However, in-depth characterization of the Ar-mTBI model is warranted to fully appreciate the diffuse pathology and range of potential behavioral effects generated by this combination of mild TBI models as compared to the repetitive linear r-mTBI model [19] and the rotational CHIMERA model [24, 25].

## Conclusions

Our findings that relatively mild but unpredictable and prolonged stress, with or without head injury, can produce long term sleep-wake dysfunction indicates the importance of the stress exposures experienced in settings with risk of repetitive head injury. Inability to maintain wakefulness can impact daily activities, such as work performance and driving safety. For service members, impaired ability to maintain wakefulness in a deployed setting could have heightened safety concerns. Our results support assessment and treatment of anxiety after TBI, which may reduce the likelihood of sleep disorders, particularly hypersomnia from excessive daytime sleepiness [31]. Furthermore, non-invasive objective assessments of daytime sleep-wake function, such as the Maintenance of Wakefulness Test (MST) and Multiple Sleep Latency Test (MSLT), may be warranted [42]. Such assessments will also support the goal of developing treatments designed for the specific pattern of sleep-wake symptoms experienced by a given patient. Our findings also emphasize the need to consider patient-specific factors, including genetics with sex/gender effects, as significant modifiers of long term outcomes.

## Acknowledgments

The authors thank Mr. Tuan Le for technical assistance. This work was initiated as a strategic concept conveyed by Dr. David L. Brody (Uniformed Services University), during his tenure as Director of the Center for Neuroscience and Regenerative Medicine (CNRM), for development of translational approaches to understand and advance treatments for the long term symptoms of military service members. We thank the program management team of the CNRM for support of these research activities.

## Disclaimer Statement

The views, information or content, and conclusions presented do not necessarily represent the official position or policy of, nor should any official endorsement be inferred on the part of, the Uniformed Services University, the Department of Defense, the U.S. Government or the Henry M. Jackson Foundation for the Advancement of Military Medicine, Inc.

## Funding Statement

These studies were funded by the United States Department of Defense within the Center for Neuroscience and Regenerative Medicine at the Uniformed Services University of the Health Sciences. The authors have no financial conflicts to disclose.

## Supplemental Materials

### STRESSORS for Unpredictable Chronic Mild Stress (UCMS)

Each stressor has a “checklist” that is filled out each time a stressor is run within the UCMS protocol.

#### Restraint

Each mouse is placed head first in a 50 ml conical Falcon plastic tube with holes at the extremity and in the cap to ensure sufficient airflow and prevent breathing concerns. The tubes are placed on a Helix 150 orbital shaker (with the shaker off) so that they do not roll around. Duration of restraint is 30 min.

#### Foot shock

Use the device in the Ugo Basile chamber. Each mouse receives 4 shocks, spaced by a random setting, over a 10 min period. Each shock is 5 sec in duration. The amplitude of each shock is 0.4 mA.

#### Rotating Shaker

Helix 150 Orbital shakers are set on a flat surface and are connected to two power strips with one toggle each. The shakers are set to 80 for speed. Two cages are placed on each shaker. Shakers are started and the cages are left on the shakers for 30 min.

#### Forced Bath

An empty rat cage (~44.5 cm long x ~23 cm wide x ~15 cm tall) is filled with 2100 mL of water (22-23 degrees Celsius). This volume of water fills the rat cage with ~2 cm of water. A mouse is placed into the rat cage and is left for 30 min.

#### Wet Bedding

A mouse cage (~26cm long x ~16cm wide x ~13cm high) is filled with 400 mL of bedding (Beta Chips, item # NOR301-62L, by ASAP; *previously used Teklad Laboratory Grade Sani-Chips 7090 by Envigo*). 250 mL of water is poured onto the bedding. The mouse is placed in the wet bedding cage sometime after 4PM and left on until the morning.

#### No Bedding

Remove bedding from cage.

#### Stroboscope

A Roxant PULSE LED strobe light is used in Standard Mode and at a low intensity of 10 Hz. Each strobe is positioned on a shelf so that each cage on the shelf is exposed to the same amount of flashing light. Strobe lights are turned on after 4PM and left on until the morning.

#### Titled Cage Platform

White wire racks (OrganizeIT wire shelving 12” deep x 36” long) are placed to lean against Helix 150 orbital shakers at a 25 degree angle. Mouse cages are placed on the tilted shelves to where the short side of the cages are at the top/bottom. The water bottle spout faces uphill. The bedding level is kept low so that it is not likely to touch the water bottle spout. The cages are placed on the tilted shelves after 4PM and left on until the morning.

#### Sleep Disruption

Near silent motorized bar sweeps across home cage floor (~9 sec) with return movement in opposite direction after 2 min delay. Mice are freely moving between the two intervals. If mouse is asleep, the sweeper motion elicits arousal due to the brief tactile stimulation. If the commonly used levels of stressors do not increase corticosterone or do not result in depressive behaviors, then the stress paradigm will be enhanced by using sleep disruption chambers between the morning and afternoon stressors.

Sleep Fragmentation Chambers (Model 80391, Lafayette Instrument) are located in B-2019. Mice are brought from LAM to B-2019 and placed in the chambers for 12 hour sessions started between 6:00-8:00 AM with the overhead lights ON. Mice are returned to LAM immediately after the 12 hr sleep disruption stressor.

Fill Sleep Fragmentation Chamber water bottles using filtered tap water in LAM G135.

Put standard mouse chow in each Sleep Fragmentation Chamber hopper.

Put 400 mL bedding (Beta Chips, item # NOR301-62L, by ASAP located in LAM; *previously used Teklad Laboratory Grade Sani-Chips 7090 by Envigo*) in each chamber.

Bring mice from G-096 to B-2019.

Put one mouse in each chamber.

Turn the cycle time selector switch to 2 min for each chamber.

Turn the power on by pressing the power button on each chamber.

Cover the light switch in B-2019 by taping a piece of paper that reads “Please do not turn off lights (initials, date)” so security or other lab personnel do not turn the lights off during the stressor.

Completely cover the door window for B-2019 by taping paper over the window.

12 hours later: turn the power off on each chamber, place each mouse back in its home cage, and return all mice to LAM.

Clean each chamber thoroughly. The chamber and cover including all attachments (sweep arm carriage, sweeper arm, guide bar and support, feeder wall, food and water support) may be submerged in water. Clean the drive mechanism, base, and stand by hand with a cloth moistened with water, mild soap solution, or disinfectant).

### BEHAVIORAL ASSESSMENTS for anxiety, depression and sleep-wake profile

#### Mouse identification

After the 3 or more days as an acclimation period, ear tags are given to mice for specific numbers. Tails are also marked to match cage assignment.

#### Weights

Immediately after the 3 or more day acclimation period, mice are weight matched to balance sham/stressed body weights. Mice that are more than 2SD above or below expected weights for C57BL/6J (see Jax data for 000664 mice) are excluded.

Weights are collected each weekday morning inside of a functioning hood. Sham mice are weighed first, then stressed mice are weighed. Stressed mice are weighed before performing any stressor procedures. Cages are never opened unless inside of a functioning hood. Clean hood thoroughly after each use.

#### Housing

Upon arrival (day 1 or day 2), housing conditions in each cage are checked to ensure that ground-dwelling enrichment is being used (tunnels or domes). If any other enrichment, such as a hanging hut, is in a cage, it will be removed and replaced with the ground-dwelling enrichment. Stressed mice do not receive any enrichment. All mice receive 3 or more days of acclimation before being handled, receiving ear tags and tail markings, and weighed. Stressed mice are then individually housed in a separate room for stressors. After completion of the stressors, mice are moved back to the regular housing room on individual ventilated cage racks and placed on different shelves from the shams. Cages are also placed so that stressed mice and sham mice do not see each other.

Sham mice receive enrichment tubes and nestlets, daily handling, and group housing whenever possible. Sham mice are group housed until after the EZM assessment, and then separated for SPT. After SPT, shams are grouped in original cages. Each cage is monitored for fighting.

Stressed mice do not receive any enrichment tubes or nestlets and are single housed throughout the study.

Tail markings for all animals are reapplied as needed to ensure animal IDs can be visualized.

Rodent bedding thru May 2020: Envigo Teklad Laboratory Grade Sani-Chips 7090. Switch to ASAP Beta Chips (item # NOR301-62L) in June 2020.

Rodent chow thru May 2020: Envigo 2018 Tekland 18% Protein Rodent Diet. Switch to ASAP PicoLab Verified – 75 IF (item # RHI5V75C3N) in June 2020.

#### Elevated Zero Maze with AnyMaze software (Stoelting, Wood Dale, IL)

Acclimation to test room:

Mice are handled calmly and quietly throughout movement, acclimation, and testing.

Sham mice and stressed mice are brought into the behavioral suite at separate times (shams first). Each cohort is brought to the behavioral suite an hour in advance to acclimate in the EZM testing room.

Lighting at low lux:

Turn off overhead lights. Turn on the two floor lights. Position the two floor lights to where they are evenly spaced between the middle of the apparatus. The floor lights should face out toward the walls on both sides. Using a light meter, the lighting is set to a low level (5-10 lux).

The elevated zero maze is used to evaluate anxiety-like behaviors. It is a circular apparatus (diameter 50 cm) divided into four equal quadrants. Two of the quadrants (located on opposite sides of the maze) are enclosed by high walls; the other two quadrants are open with 1 cm walls. The test consists of a single trial. Mice are individually placed at the edge of one of the enclosed sections of the maze, facing the inside of the closed quadrant. Each mouse is allowed to explore the maze for 5 minutes before being returned to the home cage. The time spent in each quadrant is registered from video recordings by the Any-Maze tracking software, and the time spent in the enclosed quadrants is compared to the time spent in the open quadrants, with an increased amount of time in the enclosed quadrants indicating a higher level of anxiety.

#### Sucrose Preference Test

Reagents:

Sucrose (Sigma, catalog number S3929)

Water (filtered tap water from LAM G135)

Cages (IVC with two bottle slots; Allentown)

Bottles (Stoelting, 67001-WB) with the same length spouts in each cage.

Scale for weighing bottles (analytical scale)

Complete SPT Form (Excel) for recording bottle weights.

Make sucrose solution fresh each day.

### HABITUATION and BASELINE

#### DAY 1

For the habituation phase, two bottles are prepared. Bottles are labeled “sucrose” and “water” to avoid mixing up the solutions when switching the bottles. During the afternoon of day 1, bottles with fresh 1% sucrose solution and filtered tap water are prepared and then weighed. Values are recorded onto the SPT form. Place the bottles in the testing room to sit for one hour before placing them on cages. At 3:30PM, mice are given one bottle of water (left side) and one bottle of 1% sucrose (right side). Sham mice are single housed immediately before the bottles are placed on the cages (stressed mice are already single housed).

#### DAY 2

MORNING: Bottles are weighed at 9:30AM and values are recorded onto the SPT form. Sham mice are group housed (with their original cage mates) immediately after the bottles are taken off the cages (stressed mice remain single housed). Fresh 1% sucrose is prepared during the day.

AFTERNOON: Two bottles are prepared. Bottles are labeled “sucrose” and “water” to avoid mixing up the solutions when switching the bottles. During the afternoon of day 2, bottles with fresh 1% sucrose solution and filtered tap water are prepared and then weighed. Values are recorded onto the SPT form. Place the bottles in the testing room to sit for one hour before placing them on cages. At 3:30PM, mice are given one bottle of water (right side) and one bottle of 1% sucrose (left side). Sham mice are single housed immediately before the bottles are placed on the cages (stressed mice are already single housed).

#### DAY 3

MORNING: Bottles are weighed at 9:30AM and values are recorded onto the SPT form. Sham mice are group housed (with their original cage mates) immediately after the bottles are taken off the cages (stressed mice remain single housed). Fresh 4% sucrose is prepared during the day.

AFTERNOON: Two bottles are prepared. Bottles are labeled “sucrose” and “water” to avoid mixing up the solutions when switching the bottles. During the afternoon of day 3, bottles with fresh 4% sucrose solution and filtered tap water are prepared and then weighed. Values are recorded onto the SPT form. Place the bottles in the testing room to sit for one hour before placing them on cages. At 3:30PM, mice are given one bottle of water (left side) and one bottle of 4% sucrose (right side). Sham mice are single housed immediately before the bottles are placed on the cages (stressed mice are already single housed).

#### DAY 4

MORNING: Bottles are weighed at 9:30AM and values are recorded onto the SPT form. Sham mice are group housed (with their original cage mates) immediately after the bottles are taken off the cages (stressed mice remain single housed). Fresh 0.5% sucrose is prepared during the day.

AFTERNOON: Two bottles are prepared. Bottles are labeled “sucrose” and “water” to avoid mixing up the solutions when switching the bottles. During the afternoon of day 4, bottles with fresh 0.5% sucrose solution and filtered tap water are prepared and then weighed. Values are recorded onto the SPT form. Place the bottles in the testing room to sit for one hour before placing them on cages. At 3:30PM, mice are given one bottle of water (right side) and one bottle of 0.5% sucrose (left side). Sham mice are single housed immediately before the bottles are placed on the cages (stressed mice are already single housed).

#### DAY 5

MORNING: Bottles are weighed at 9:30AM and values are recorded onto the SPT form. Sham mice are group housed (with their original cage mates) immediately after the bottles are taken off (stressed mice remain single housed).

Mice that have a sucrose preference of <60% during habituation are determined to be non-responsive and are omitted from the data analysis.

### TESTING

#### DAY 1

For the testing phase, two bottles are prepared. Bottles are labeled “sucrose” and “water” to avoid mixing up the solutions when switching the bottles. During the afternoon of day 1, bottles with fresh 0.5% sucrose solution and filtered tap water are prepared and then weighed. Values are recorded onto the SPT form. Place the bottles in the testing room to sit for one hour before placing them on the cages. At 3:30PM, half of the mice from each group are given one bottle of water on the left side and one bottle of 0.5% sucrose on the right side, and the other half of the mice from each group are given one bottle of water on the right side and one bottle of 0.5% sucrose on the left side. Sham mice are single housed immediately before the bottles are placed on the cages (stressed mice are already single housed).

#### DAY 2

Bottles are weighed at 9:30AM and values are recorded onto the SPT form. Sham mice are group housed (with their original cage mates) immediately after the bottles are taken off the cages (stressed mice remain single housed).

Bottle weights are recorded with 3 digits beyond the decimal point using an analytical scale that measures to 4 digits (0.0000 g). Total consumption of sucrose and water is measured in grams (for example, 4.394 g). The percentage of sucrose preference per day is calculated as follows: (consumed sucrose*100)/(consumed water + consumed sucrose).

#### PROCESS FOR BOTTLE PLACEMENT

- Bottles are first labeled either “sucrose” or “water.”
- All sucrose bottles are prepared and placed in one Styrofoam holder, and all water bottles are prepared and placed in a second (separate) Styrofoam holder.
- Bottles are placed in the testing room one hour before being placed on cages.
- One sucrose bottle is weighed and placed on each mouse cage (the side of the cage depends on the day of habituation or testing). All sucrose bottles are weighed and placed on the cages before any water bottle is placed on a cage.
- One water bottle is weighed and placed on each mouse cage (the side of the cage depends on the day of habituation or testing).
- Each cage is checked to ensure that there is one sucrose bottle and one water bottle in the correct locations for each cage.

#### PROCESS FOR BOTTLE REMOVAL

- Each sucrose bottle is removed and weighed. One bottle should be removed and (immediately) weighed at a time. All sucrose bottles are removed before any water bottle is removed from a cage.
- Each water bottle is removed and weighed. One bottle should be removed and (immediately) weighed at a time.

#### Tail Suspension (Bioseb In Vivo Research Instruments, bioseb.com)

Acclimation to test room:

Handle mice calmly and quietly throughout movement, acclimation, and testing.

Sham mice and stressed mice are brought into the behavioral suite at separate times (shams first). Each cohort is brought to the behavioral suite an hour in advance to acclimate in the TST testing room.

Lighting at low lux:

Turn off overhead lights. Turn on the single floor light in the room. The floor light should face out toward the wall. Using a light meter, the lighting is set to a low level (5-10 lux).

Immobility of a mouse suspended by the tail is timed as a measure of behavioral despair. The mouse is placed in an automated tail suspension apparatus designed to measure the mouse movement and record the immobility time (Bioseb). The mice are weighed and placed in the experimental room at least 60 min prior to testing. To ensure that the mice hang downward and cannot climb up to any parts of the apparatus or tape, the tail is passed through a 4-cm length of hollow polycarbonate tubing (1.3 cm inner diameter;McMaster-Carr, Santa Fe Springs, CA; #8585K41; Can et al. 2012 PMID 22315011). A piece of adhesive tape is wrapped around the tail in a constant position of approximately three quarters of the distance from the base of the tail.

The suspension hook is passed through the adhesive within 1-2 mm of the tail. The mouse is continuously observed for 6 min during automated recording of the immobility time. Mice will initially exhibit vigorous movement and become immobile after a few minutes.

#### Social Interaction (3-chamber from Ugo Basile, Italy; with AnyMaze software, Stoelting, Wood Dale, IL)

Handle mice calmly and quietly throughout movement, acclimation and testing.

Sham mice and stressed mice are brought into the behavioral suite at separate times (shams first). Each cohort is brought to the behavioral suite an hour in advance to acclimate. NOTE that the experimental cohort for the social assay is NOT put into the testing room for acclimation or during testing because the social testing can be influenced by sounds from the mice that can distract the experimental mouse from interacting with the mouse in the carrier(s).The cohort of experimental mice acclimates in a nearby quiet and empty room. The experimental mice are kept in that same room while waiting to be tested. This room should have lighting set up similar to the testing room (see below) – red light on and an extra floor light on for overall dim lighting.

At the time of testing, one experimental mouse is transferred to a clean empty cage with bedding, so 12 extra cages will need to be brought to the acclimation room if testing 12 experimental mice. Cover this cage with something clean to block the experimental mouse from the bright hallway lighting when carrying from the acclimation room to the experimental room. Bring the experimental mouse back to the acclimation room the same way after testing.

Mice are placed in the 3 chamber sociability apparatus (Stoelting; Wood Dale, IL) for testing social behavior.

Test room lighting as low lux:

Turn off overhead lights. Turn on the two floor lights. Position the two floor lights to where they are evenly spaced between the middle chamber of the apparatus. The floor light should face out toward the walls on both sides. Using a light meter, the lighting is set to approximately 15 lux. Two light meter readings should be taken: one from the top of each enclosure.

AnyMaze:

Open AnyMaze

Create a new experiment

Name the trial and save the experiment

Select “protocol”

Open “TTC two camera”

Add testing mice data (mouse ID, treatment group, zone)

*the social zone (stranger #1) and the empty/novel zone (stranger #2) should be counterbalanced, alternating the “top” and “bottom” chambers of the apparatus for every testing animal*

The two cameras need to be aligned so that the apparatus chambers are in line with the cameras each time a new experiment is ran.

There are four trials per mouse. All trials are performed for each mouse before moving on to the next mouse. Before the beginning of each of the four trials, it is also extremely important that the experimenter perform a visual check on the computer to ensure that both of the wired enclosures are centered under the cameras (the wired enclosures must align so that the view of the carrier does not show the bottom ring when properly blocked by the upper parts of the wired enclosure). It is also extremely important that the experimenter perform a visual check to ensure both carriers are concentric within each scoring circle before the start of each trial.

TRIAL 1: (Start with both plastic doors in) The first 5-min session is an open habituation session allowing the experimental mouse free access to all three chambers. There are no wired enclosures in the apparatus during this trial. Place the testing mouse in the center chamber. Click start on AnyMaze. Immediately remove both plastic doors so the testing mouse is allowed free access to all three chambers. Verify that the mouse is being tracked (orange dot on black) and leave the room. When trial 1 is complete, guide the testing mouse to the center of the apparatus and place the plastic doors back on.

TRIAL 2: The second 5-min session is another habituation session in which wired enclosures are placed in the two outer chambers, and the testing mouse is allowed free access to all three chambers. Before the trial, place the two wired enclosures in the outer chambers and ensure they are centered with orange outlines on the software. Place the testing mouse in the center chamber. Click start on AnyMaze. Immediately remove both plastic doors so the testing mouse is allowed free access to all three chambers. Verify that the mouse is being tracked (orange dot on black) and leave the room. When trial 2 is complete, guide the testing mouse to the center of the apparatus and place the plastic doors back on.

TRIAL 3 (social approach): The third trial is a 10-min trial in which a gender-matched conspecific mouse (#1) is introduced into one of the two wire enclosures within the apparatus. Place conspecific mouse (#1) into the enclosure, either “top” or “bottom.” ‘Top” or “bottom” location will change between testing mice, and the software will tell you which one you should use. Place the testing mouse in the center chamber. Click start on AnyMaze. Immediately remove both plastic doors so the testing mouse is allowed free access to all three chambers. Verify that the testing mouse is being tracked (orange dot on black) and leave the room. When trial 3 is complete, guide the testing mouse to the center of the apparatus and place the plastic doors back on. Leave the conspecific mouse (#1) in the wire enclosure.

TRIAL 4 (social memory): The fourth 10-min trial is a social novelty trial in which a second gender-matched conspecific mouse (#2) is introduced in the remaining wire enclosure within the apparatus. Place conspecific mouse (#2) into the remaining enclosure. Place the testing mouse in the center chamber. Click start on AnyMaze. Immediately remove both plastic doors so the testing mouse is allowed free access to all three chambers. Verify that the testing mouse is being tracked (orange dot on black) and leave the room. The social behavior test is complete at the end of trial 4. After the mice are removed from the chamber, clean the chamber and both enclosures thoroughly with 70% EtOH.

All sessions are video recorded to facilitate scoring of the time spent within each of the 3 chambers. Video files are stored and used in accordance with IACUC policy. The video files are used for deriving quantitative values for data analysis. The quantitative data is used in publications and presentations. The video files are not used in publications, presentations, or teaching.

Exclusion criteria: If sham mice do not show a preference for the intruder mouse vs the empty carrier during trial 3 (social approach), both the social approach trial and the social memory trial are determined to be invalid. If sham mice show a preference for conspecific mouse (#1) vs the empty carrier during trial 3 (social approach) but do not show a preference for the familiar conspecific mouse (#1) vs the unfamiliar conspecific mouse (#2) during trial 4 (social memory), the social memory test is determined to be invalid.

#### CONSPECIFIC MICE

Training:

Conspecific mice are habituated to the wire enclosures for 15 min daily for 5 consecutive days before the sociability test. Each intruder is placed in an enclosure for 15 min daily (only one conspecific mouse is trained at a time), and the experimenter leaves the room and observes through the door window each conspecific’s climbing behavior daily. Any atypical or excessive climbing behaviors are recorded each day on the Training Form. At the end of the 5 day training period, any mouse that has been determined to have excessive climbing behavior or other atypical behaviors that may interfere with the social interaction test will be excluded. The chamber and the enclosure are thoroughly cleaned with 70% EtOH after each mouse.

Intruder handling during testing:

Intruder mice are kept in the quite area outside of the experimental room while waiting. Keep conspecific mice away from noise or other experiments. Conspecific mice are used an equal amount of times and are rotated in sequential order to use for testing mice. Half of the mice should be rotated as conspecific #1, and half of the mice should be rotated as conspecific #2 for a given day. If any conspecific is observed to be climbing excessively on the enclosure or aggressive to an experimental mouse, that intruder is taken out of the rotation for the remainder of the social behavior test. The conspecific mouse IDs used with each testing mouse are recorded on the Testing Form.

#### Piezo sleep sensors (PiezoSleep Mouse Behavioral Tracking System, Signal Solutions, LLC)

The Piezo sleep sensor system is a fully automated, non-invasive system that monitors and analyzes data regarding sleep/wake cycles. The sleep wake monitoring system is based on piezoelectric sensor technology that transforms mechanical pressure into electrical signals. For in depth system details and operation, refer to instruction manuals provided by Signal Solutions.

Hardware:

12 Customized Open-Floor Cages with water bottles

Non-invasive Piezo Floor Sensors (PZ-77)

Data Acquisition Sensors (DAQ)

USB 3.0 to USB Micro Cable (24 ft)

Squid Box 16-channel System

Backup power supply

Desktop

Software:

PiezoSleep software is used for data collection

SleepStats Data Explorer software is used for data analysis

Forms to complete for experimental record:

Piezo Sleep Sensor Animal ID Form

Piezo Sleep Sensor Light Level Log

Piezo Sleep Sensor Daily Monitor Log

Piezo Sleep Sensor Sanitation Log

*This protocol is for a 72hr testing period.*

##### DAY 1 – in the morning for setup

Place “EXPERIMENT IN PROGRESS: DO NOT ENTER” sign on door of testing room.

Center cage rack on back wall to avoid air flow from vents.

Confirm each floor sensor pad is aligned with the length and width of the chamber, and a cage shield (thin plastic protector on floor) is placed between the chamber and the sensor.

Confirm the correct chamber is plugged into the intended channel on the Squid Box.

Place 0.5cm (~180mL) bedding (Teklad Laboratory Grade Sani-Chips 7090 by Envigo, located in LAM) in each chamber (no neslets or enrichment).

Fill food (standard mouse chow) and filtered water for each chamber.

Record the light level using a light meter (place the light meter front + centered on each rack; 3 levels will be recorded – 1 for each of the 3 racks).

Complete the “Piezo Sleep Sensor Animal ID Form” to record which mouse is associated with each chamber/channel #.

##### At ~11:30AM

Put one mouse in each chamber.

Turn on power supply (keep at default medium sensitivity setting), computer, and Squid Box.

Open “Piezo_Sleep2.18” by clicking on the icon.

Select “USB-6221” from the pull down menu and press “OK”.

Enter the number of active channels (i.e., how many mice being run) and press “OK”.

For “Channel Setup”, set “Dark-to-Light Time” at 06:00 and “Light-to-Dark Time” at 18:00, then enter animal IDs for each “Channel Number”.

Enter file name (Cohort #, date) to store the data, and save to the desktop folder “TTC Piezo Sleep Studies Data”.

Review your selections, and if no changes are necessary click “No re-selection needed” and press “OK” **at 12PM**.

(Experiment is now started and will run for 72 hours).

##### DAY 2

Being as quiet and brief as possible, check on mice at ~9AM and at ~5PM to ensure:

-system is running properly

-all animals have plenty of food and water

Use the “Piezo Sleep Sensors Daily Monitor Log” to record your name, date, time that you checked on the mice, and any notes such as adding food/water. *Only refill food and/or water if the levels are low.*

##### DAY 3

Being as quiet and brief as possible, check on mice at ~9AM and at ~5PM to ensure:

-system is running properly

-all animals have plenty of food and water

Use the “Piezo Sleep Sensors Daily Monitor Log” to record your name, date, time that you checked on the mice, and any notes such as adding food/water. *Only refill food and/or water if the levels are low.*

##### DAY 4 at 12PM

Stop the experiment by pressing “STOP” on Piezo Sleep.

Record the light level using a light meter.

House mice in their housing room in standard cages. Sham mice are always group housed with their original cage mates.

CLEANING: refer to “Piezo Sleep-Wake Single-Cage System Assembly” manual.

Complete the “Piezo Sleep Sensor Sanitation Log”.

## Notes

### Competing Interest Statement

The authors have declared no competing interest.

